# Cryo-EM structure of the naked mole-rat ribosome reveals a stabilized split 28S rRNA

**DOI:** 10.1101/2025.10.26.684699

**Authors:** Mehmet Gül, Alice Rossi, Christian M.T. Spahn, Gary R. Lewin, Mikhail Kudryashev

## Abstract

The naked mole-rat *(Heterocephalus glaber)* is a long-lived mammal with remarkable resistance to cancer and hypoxia, suggesting the evolution of robust proteostasis networks. The ribosome, the central engine of protein synthesis, is key for cellular stress responses and has an unusual feature of the 28S rRNA split; however, the details of its organization remain unknown. Here, we present high-resolution cryo-EM structures of the naked mole-rat 80S ribosome in two states of the elongation cycle. The structures reveal a conserved overall architecture and rRNA modification landscape compared to other mammals, and provide an atomic-level view of the unique break in the 28S rRNA. This cleavage event, located in the D6 expansion segment, is structurally stabilized by a network of interactions with surrounding ribosomal proteins, maintaining the integrity of the large subunit. Our comparative analysis revealed that this compensatory network preserves a canonical architecture nearly indistinguishable from intact mouse and human ribosomes. These findings resolve the structural basis of this unique cleavage, showing that it is a stable, integrated feature whose function is likely linked to more subtle regulatory mechanisms, rather than inducing major structural rearrangements.

## INTRODUCTION

The naked mole-rat (NMR) is a subterranean rodent with slower than typical mammalian aging, exhibiting extraordinary longevity, sustained fertility, and remarkable resistance to age-related diseases, including cancer and neurodegeneration^1,2^. This unique biology is coupled with tolerance to extreme environments, particularly severe hypoxia, a condition that would be lethal to most other mammals^3,4^. These physiological traits suggest the existence of highly robust cellular maintenance programs, particularly the network of pathways that maintain protein homeostasis, or proteostasis ^5^. The ribosome, the universally conserved ribonucleoprotein complex responsible for protein synthesis, is central to proteostasis. It is a major consumer of cellular energy and a primary control point for managing cellular resources, especially during periods of environmental stress, such as hypoxia, where rapid translational repression is a critical survival strategy. Ribosomes from many species have been extensively studied by structural methods^6–8^, including the detailed understanding of their regulation^9,10^; however, structural insights into the ribosomes of NMRs are missing.

Previous work has provided insights that the NMR translation machinery may possess unique characteristics. Fibroblasts from NMRs exhibit increased translational fidelity compared to those from mice, and, most strikingly, a unique processing event was identified in the 28S ribosomal RNA (rRNA)^5^. This event, termed a “hidden break,” results from the excision of a ∼260-nucleotide fragment from the D6 expansion segment during rRNA maturation, leaving two fragments that remain associated within the large ribosomal subunit. This raises a question: how is the structural and functional integrity of the large ribosomal subunit maintained despite a break in its rRNA backbone? The structural basis and functional consequences of this unusual rRNA architecture have remained unknown.

Comparative studies support the view that ribosomal heterogeneity can have functional consequences. In *Plasmodium* spp., for example, the presence of distinct stage-specific rRNA alleles with divergent expansion segments enables specialized translation programs during the parasite life cycle^11,12^. While mammalian rDNA is generally homogenized across tandem repeats, subtle sequence and structural variations do exist and may impact translation efficiency and regulation^12^. A high-resolution comparison of naked mole-rat ribosomes with those of closely related mammals is therefore essential for distinguishing universally conserved features from potential adaptations.

To address this, we used single-particle cryo-EM to determine the structure of the NMR ribosome across multiple functional conformations, enabling a rigorous, quantitative comparison with other mammalian ribosomes^6,13,14^. In this study, we determined the structures of the NMR 80S ribosome purified from NMR liver. We provide an atomic-level visualization of the NMR ribosome in two distinct states and describe the break within the 28S rRNA. Our comparative analysis reveals that while the overall ribosome structure is highly conserved compared to those of other mammals, the cleavage site is a stable, fully integrated feature. A conserved scaffold of ribosomal proteins maintains the rRNA fragments in a canonical conformation, preserving the energetic stability of the region. Our work thus defines the structural basis of a unique, species-specific feature of the NMR ribosome, demonstrating that it is a stable element rather than a sign of instability.

## RESULTS

### Cryo-EM structures of the naked mole-rat 80S ribosome capture two states of the elongation cycle

To investigate the structural basis of protein synthesis in the NMR, 80S ribosomes were purified from liver tissue and subjected to single-particle cryo-EM. The analysis resulted in two predominant structurally distinct conformational states, representing the intermediates of the translation elongation cycle, a post-translocation (POST) state with tRNAs in the P and E sites, and a rotated-2 pre-translocation state (rotated-2 PRE) with a rotated SSU and hybrid A/P and P/E tRNAs. The final reconstructions had resolutions of 2.9 Å for the POST state and 2.7 Å for the rotated-2 PRE state (Figure 1A-B, Suppl Figure S1, S2, Table S1). The quality of the cryo-EM density maps is high throughout the conserved core of both the large 60S (LSU) and small 40S (SSU) subunits, with well-resolved rRNA bases and protein side chains that enabled confident and accurate atomic model building (see Methods).

**Figure 1.**
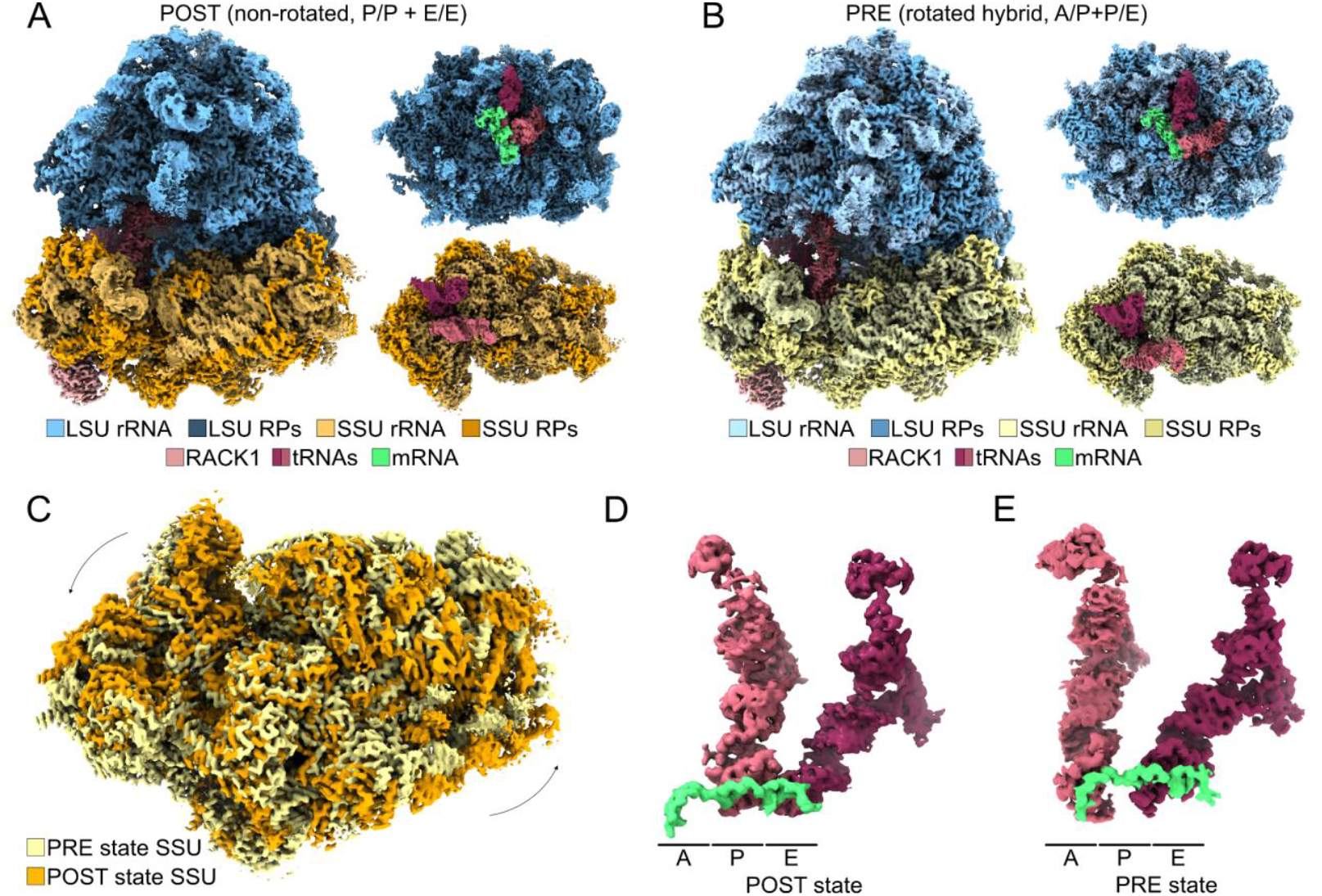
**Cryo-EM density maps of the naked mole-rat 80S ribosome** in (A) the POST non-rotated, P/P + E/E state and (B) the PRE, rotated hybrid, A/P+P/E state. In both panels, the 60S rRNA is shown in lighter blue and the 60S ribosomal proteins in darker blue, while the 40S rRNA is shown in lighter yellow and the 40S ribosomal proteins in orange (POST) and darker yellow (PRE). The tRNA in the P site (non-rotated state) and A/P hybrid site (rotated state) is shown in dark pink, the tRNA in the E site (non-rotated) and P/E site (rotated) in light pink, and the mRNA in green. (C) Superposition of the 40S subunits, highlighting the counterclockwise rotation of the body, is indicated by black arrows. (D) tRNA and mRNA densities in the POST state, with tRNAs in dark pink and light pink, and mRNA in green. A, P, and E-sites are indicated. (E) tRNA and mRNA densities in the rotated-2 PRE state, with hybrid-state tRNAs in dark and light pink and mRNA in green.

The two states differ primarily by a rigid-body rotation of the SSU relative to the LSU. Superposition of the two models based on the 60S subunit revealed a counter-clockwise rotation of the 40S subunit by approximately 11^°^ (Figure 1C). These global conformational changes represent the canonical transition between classical and rotated states of the ribosome during elongation. Consistent with their assignment as distinct functional intermediates, the two structures also exhibit different configurations of bound tRNAs. In the POST state, the density maps show tRNAs occupying the classical peptidyl (P) and exit (E) sites (Figure 1D). In contrast, the rotated-2 PRE state captures tRNAs in A/P and P/E hybrid configurations, where the acceptor ends of the tRNAs have translocated on the 60S subunit while their anticodon stems remain anchored to the 40S subunit (Figure 1E). The capture of these canonical, physiologically relevant conformations confirms the functional integrity of the purified ribosomes. It provides a framework for a detailed comparative analysis of the naked mole-rat translation machinery.

Collectively, these observations indicate that the PRE and POST states captured in the naked mole-rat ribosome represent distinct elongation intermediates, differing not only in the relative orientation of the ribosomal subunits but also in the positions of bound tRNAs. The structural features observed here align with conformational changes observed in the elongation mechanism in eukaryotes^15,16^.

### The naked mole-rat ribosome shares a highly conserved architecture with its mammalian counterparts

To identify possible species-specific features from the universally conserved architecture of the mammalian ribosomes, we compared the structure of the PRE state NMR ribosome with the structures from the mouse (*Mus musculus*) kidney (PDB: 7CPU)^17^ and human HeLa cells (PDB: 6QZP)^7^, both resolved to a similar global resolution. To eliminate the influence of possible different states, the superpositions were performed for the subunits individually (Figure 2A, 2B). In all comparisons, the conserved cores of the LSU and SSU remain uniformly low in deviation with lower RMSD values, where the solvent-exposed loops and flexible loops on the intersubunit face show higher differences. Our analysis using the high-resolution human structure reveals an even higher degree of conservation than previously appreciated. For the LSU, deviations in the NMR–human comparison are exceptionally low, confirming a near-identical core architecture. Similarly, the SSU shows strong conservation with minimal deviations. Residue-wise analysis of the 28S rRNA (Figure 2C) now shows that the NMR–human comparison (orange) exhibits a remarkably low baseline deviation, with discrete peaks corresponding primarily to the most flexible, solvent-exposed expansion segments. The comparison with the mouse structure (blue) remains consistent, showing low overall deviation.

**Figure 2.**
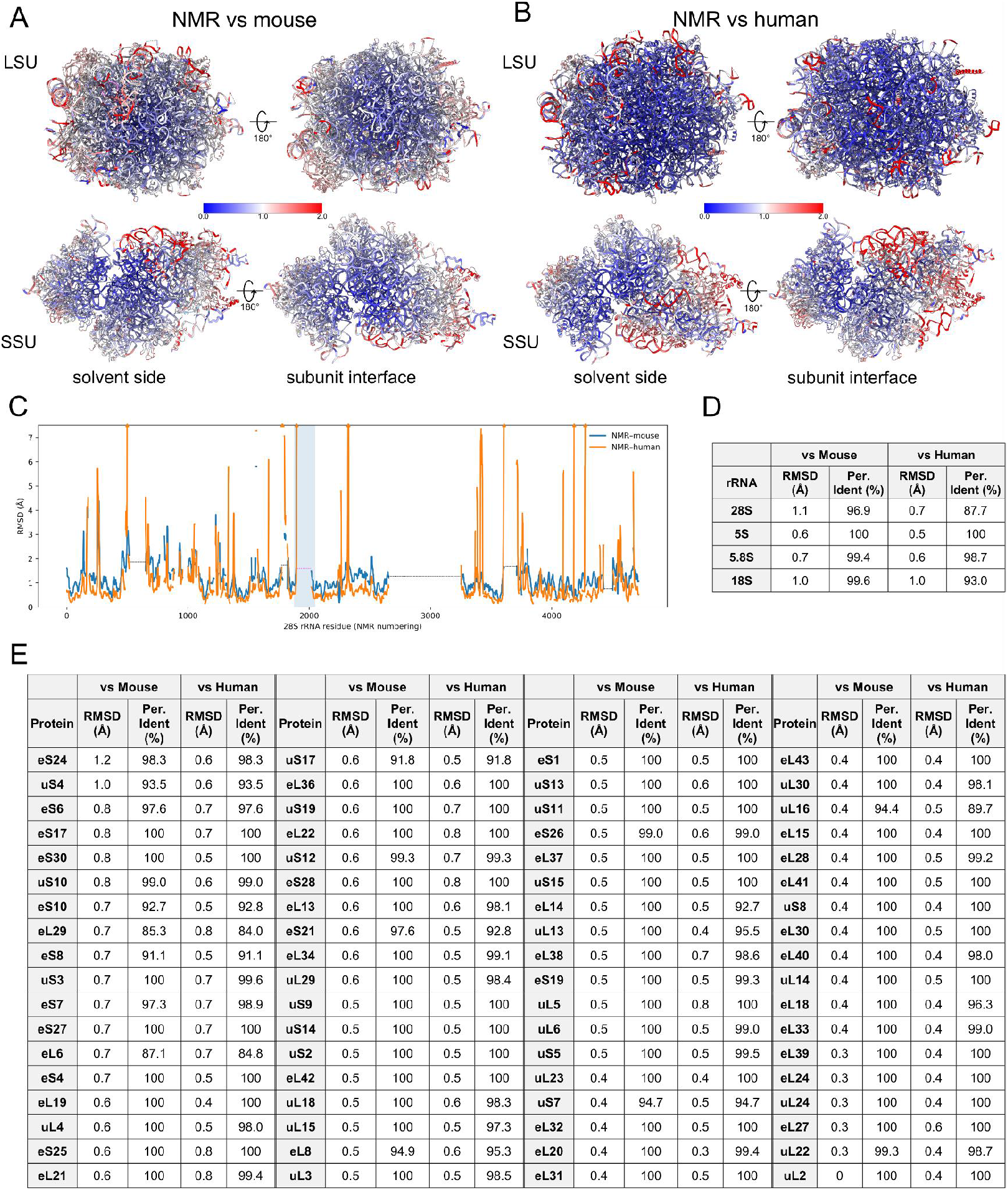
Structural comparison of the naked mole-rat and mouse 80S ribosomes in the rotated state. (A) Superposition of the naked mole-rat ribosome and the mouse ribosome subunits, colored by Cα RMSD values. (B) Superposition of the naked mole-rat ribosome and the human ribosome subunits colored by Cα RMSD values. (C) RMSD-per-residue plot of the naked mole-rat 28S rRNA with mouse (blue) and human (orange). The unmodelled gaps in the chain are indicated with dashed lines. The 28S rRNA cleavage site is highlighted with a pale blue background. (D) Phosphate backbone RMSD analysis and sequence identity of rRNAs between NMR and mouse/human. (D) Cα RMSD and sequence identity comparison of ribosomal proteins between NMR and mouse/human.

The majority of ribosomal proteins exhibit high sequence identity and low Cα RMSD values when compared to both mouse and human orthologs, indicating strong structural homology (Figure 2E). Deviations are largely confined to the solvent-exposed termini and flexible loop regions of peripheral proteins, consistent with known patterns of interspecies variability^18^ (Figure 2A). Similarly, analysis of the rRNAs shows that the sequence identities and the core secondary and tertiary structures are nearly identical, with the most significant deviations localized to the tips of several eukaryotic-specific rRNA expansion segments (ESs) (Figure 2D). This high degree of overall structural conservation underscores the strong evolutionary pressure to maintain the core functional machinery of translation. Within this largely conserved framework, one region stands out as a dramatic and unique structural divergence specific to the NMR.

### The 28S rRNA cleavage is a stable feature that does not perturb the local protein environment

A key point of comparison is the D6 region of the 28S rRNA, where a 28S rRNA cleavage site has been previously reported in naked mole-rats^5^. Focused examination of the 28S rRNA D6 region revealed a structural discontinuity in the NMR ribosome, consistent with the reported cleavage (Figure 3A-D). In the cryo-EM maps, density could be traced on both sides of the cleavage site, with no continuous density connecting the two fragments. By contrast, the equivalent region in the mouse ribosome has continuous but fragmented density. This lower local resolution limits the precision of rRNA modeling but still supports an intact 28S rRNA configuration. This comparison highlights the presence of a structurally well-resolved rRNA discontinuity in the naked mole-rat ribosome, contrasted with the intact but flexible corresponding region in the mouse.

**Figure 3.**
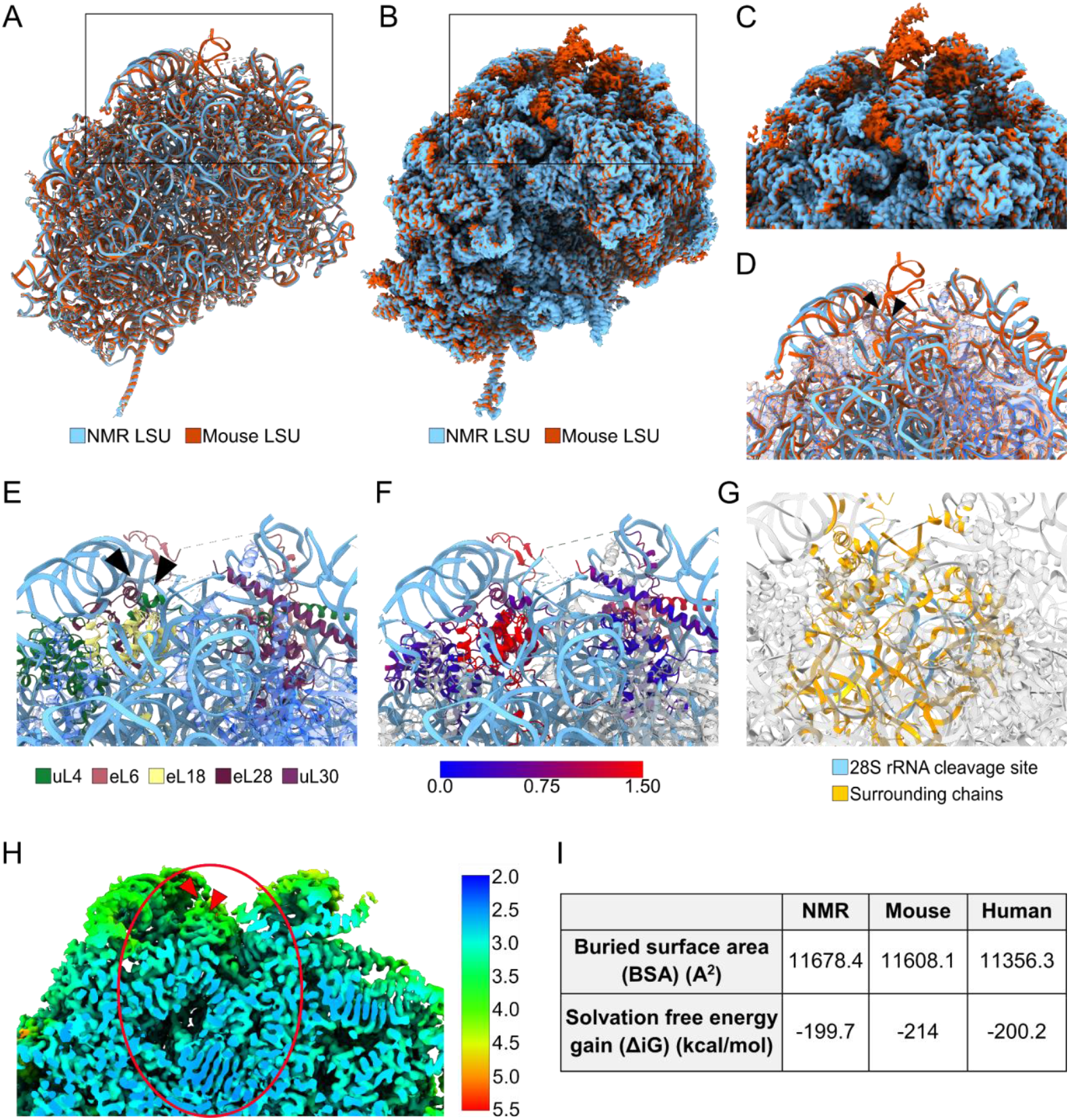
Comparison of structures of the 28S rRNA D6 region in naked mole-rat and mouse ribosomes. (A) Overall superposition of the naked mole-rat and mouse large ribosomal subunits to visualize the D6 domain of the 28S rRNA. (B) Superposition of the NMR and mouse cryo-EM maps. (C) Close-up view of the D6 cleavage site, with the naked mole-rat map shown in blue and the mouse map in red. The rRNA break points are indicated with arrowheads. (D) Close-up view of the cleavage site on the superposed atomic models of the NMR and mouse ribosomes. (E) Cleavage site in the NMR ribosome, with 28S rRNA shown in light blue and surrounding proteins in different colors. The rest of the proteins are displayed with increased transparency to improve visibility. (F) Cleavage site in the NMR ribosome, with 28S rRNA shown in light blue and surrounding proteins colored by RMSD. A scale bar for RMSD values is provided. (G) The region extracted for PDBePISA analysis is shown in light blue (28S rRNA fragment) and orange (surrounding chains). (H) Close-up view of the D6 cleavage site and its surroundings, colored by local resolution. The rRNA break points are indicated with arrowheads. The region extracted for PDBePISA analysis is annotated with a red ellipse. (I) Buried surface area and solvation free energy gain values for NMR, mouse, and human ribosomes in the NMR 28S rRNA cleavage site.

Local resolution maps of the NMR ribosome showed uniform resolution and suggested that the discontinuity in the NMR ribosome is a feature, not a technical artefact. The map density is well-resolved on both flanks of the gap, and the region as a whole is of high resolution (Figure 3H). The missing density is therefore a true rRNA break, unlike in the mouse ribosome, where this region is intact but flexible. Mapping the local environment around the cleavage site showed that several ribosomal proteins (uL4, eL6, eL18, eL28, and uL30) are positioned to support the two rRNA fragments (Figure 3E). The proteins at this site are positioned similarly in both the naked mole-rat and mouse ribosomes, with RMSDs below 1.5 Å (Figure 3F), and this arrangement is likewise preserved when compared to the high-resolution human structure. These proteins likely stabilize the break through a network of interactions, helping to maintain the integrity of the large subunit despite the discontinuity. A 28S rRNA fragment at the cleavage site and all proteins within a 12 Å radius were extracted for PDBePISA analysis (Figure 3G). Calculations of the buried surface area and solvation free energy at the interface showed that the degree of protein-RNA contacts and the energetic favorability of the interface are comparable across NMR, mouse, and human (Figure 3I). This analysis demonstrates that the protein-rRNA interface is not energetically compromised by the cleavage, providing evidence for a fully compensated and stable architecture of the ribosome.

### The naked mole-rat ribosome maintains a conserved pattern of rRNA modifications and magnesium ion coordination

Eukaryotic rRNAs are decorated with chemical modifications, primarily 2’-O-methylations (Nm) and pseudouridylations (Ψ), which play crucial roles in ribosome assembly and the fine-tuning of translational function^7,9,19,20^. The high quality of our cryo-EM maps enabled a systematic survey of the rRNA modification landscape in the NMR ribosome. By inspecting the density for features exceeding the standard RNA chemistry, we identified a total of 10 putative modification sites across the 18S, 5.8S, and 28S rRNAs (Figure 4A, Table 1). These modifications, found in conserved functional regions, reflect the epitranscriptomic signatures previously observed in mammalian ribosomes ^21^.

**Table 1:**
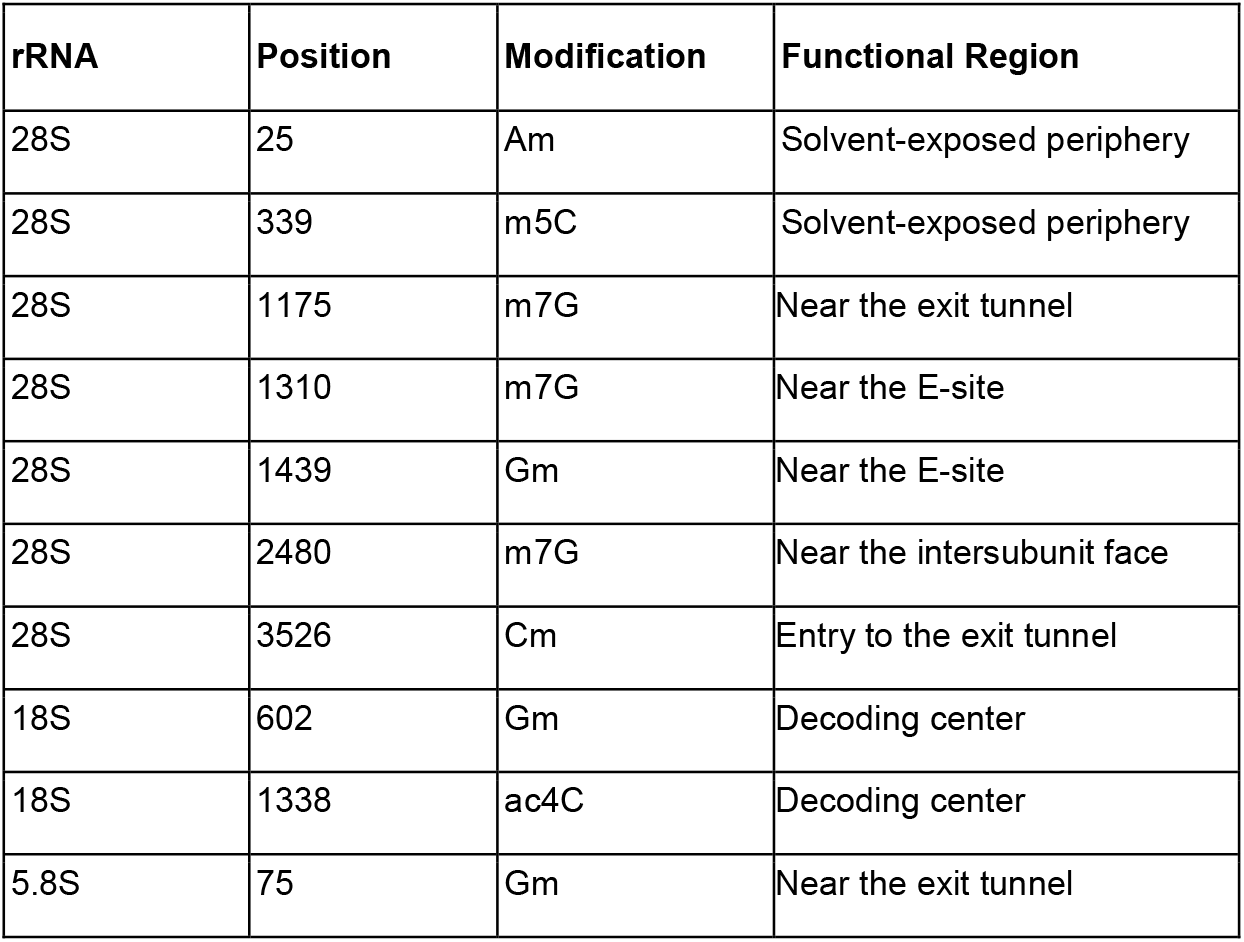
Putative rRNA Modifications Identified in the Naked Mole-Rat Ribosome.

**Figure 4.**
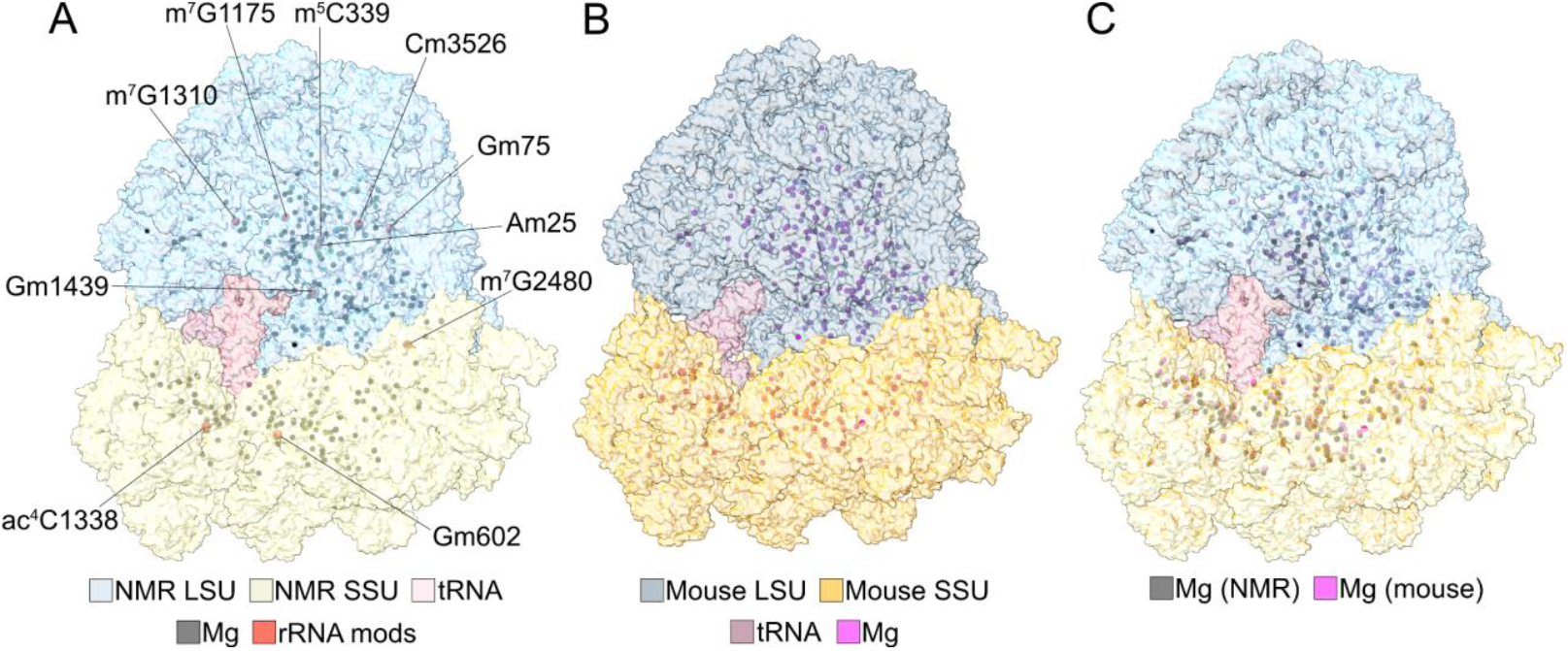
rRNA modifications and Mg^2^+ ion distribution. (A) rRNA modifications and Mg ion distribution in the naked mole-rat ribosome. (B) Mg ion distribution in the mouse ribosome. (C) Superposition of the NMR and mouse ribosomes, comparing the distribution of Mg ions

The distribution of Mg^2^+ ions in the NMR ribosome showed numerous Mg^2^+ ion clusters within the rRNA core, reflecting their role in stabilizing tertiary structure (Figure 4A). Comparison with the mouse ribosome ^22^ (Figure 4B) revealed an overall similar pattern of Mg^2^+ binding sites, and superposition of the two species showed only minor shifts in ion positions (Figure 4C). These observations suggest that the NMR ribosome maintains a conserved network of rRNA modifications and Mg^2^+ coordination sites. Together, these features confirm that despite the break in the 28S rRNA, the ribosome’s core functional centers and overall structural integrity are precisely maintained.

## DISCUSSION

In this study, we present high-resolution cryo-EM structures of the naked mole-rat 80S ribosome, capturing two distinct conformational states of the translation elongation cycle. Our analysis reveals that while the overall architecture, ribosomal protein composition, and epitranscriptomic landscape are highly conserved with other mammals, the NMR ribosome possesses a unique structural feature: a specific cleavage in the D6 region of the 28S rRNA. These structures provide atomic-level visualization of this “hidden break,” previously identified biochemically^5^, and offer a structural foundation.

Our work documents the discontinuity in the 28S rRNA, which remodels the D6 region. Unlike a site of degradation or structural instability, the cleavage site appears to be a stable, defined feature. The two resulting rRNA fragments are held in place by a conserved network of surrounding ribosomal proteins (uL4, eL6, eL18, eL28, and uL30), which show minimal positional deviation compared to their mouse and human counterparts. Furthermore, our interface analysis suggests that the protein-RNA contacts and the energetic stability of this region are comparable to those in mouse and human ribosomes, which possess an intact rRNA backbone. Our findings are consistent with a recent report by Gutierrez-Vargas and colleagues, who also observed the overall preservation of the ribosome core architecture^23^. The work presented here extends this finding by providing a higher-resolution view of two distinct functional states of the elongation cycle, a quantitative analysis of the energetic stability at the cleavage interface, and a complete map of the conserved rRNA modification landscape, together providing a detailed understanding of the functional integrity of the NMR ribosome.

A hypothesis was that this cleavage might introduce local flexibility or remodel the D6 region, potentially as a mechanism to influence translational fidelity or adapt to stress. However, our comparative structural analyses point to the contrary. Despite the discontinuity in the backbone, the local architecture is well-preserved. The surrounding ribosomal proteins maintain their canonical positions, and the energetic stability of the protein-RNA interface is not compromised. This suggests that the NMR ribosome has evolved to compensate for the break, indicating a strong selective pressure to preserve the native architecture of this region. Therefore, the biological significance of this feature must lie in a more subtle mechanism than major structural rearrangement.

In conclusion, our work provides an atomic-resolution view of the naked mole-rat ribosome, shedding light on the structural nature of its unique 28S rRNA cleavage. We showed that this break is a stable, integral feature of an otherwise highly conserved ribosome. The structure compensates for the break, maintaining the native architecture and energetic stability of the D6 region. These findings lay the groundwork for future investigations to determine whether this unique rRNA architecture plays a more subtle role in ribosome biogenesis, turnover, or the extraordinary stress resistance of this species.

## MATERIALS AND METHODS

### Purification of ribosomes from naked mole-rat liver tissue

Liver tissue was dissected from an adult naked mole-rat and immediately transferred to a chilled dish on ice. All subsequent steps were carried out under ice-cold conditions. The tissue was washed with ice-cold PBS, chopped into small pieces, and homogenized in lysis buffer (20 mM Tris-HCl, pH 7.4, 10 mM MgCl_2_, 150 mM KCl, 2 mM DTT, protease inhibitor cocktail) using glass beads and a Precellys Evolution Touch homogenizer. The homogenate was clarified by centrifugation at 10,000 × g for 10 min to remove cell debris. The supernatant was layered onto a 30% sucrose cushion and centrifuged for 2 h at 100,000 × g. The resulting pellet was resuspended in a minimal volume of wash buffer (20 mM Tris-HCl, pH 7.4, 10 mM MgCl_2_, 150 mM KCl) and further purified by sucrose density gradient centrifugation (25–50% sucrose in 5% increments) for 3 h at 100,000 × g. Fractions containing ribosomes were identified by absorbance at 254 nm using a NanoDrop spectrophotometer, pooled, pelleted, and resuspended in sucrose-free buffer (20 mM Tris-HCl, pH 7.4, 10 mM MgCl_2_, 150 mM KCl).

### Cryo-EM grid preparation

Cu300 R2/2 grids with an additional 2 nm amorphous carbon support film (Quantifoil Micro Tools) were glow-discharged in a PELCO easiGlow Glow Discharge Cleaning System at 15 mA for 45 s. 4 µl of purified ribosome sample was applied to each grid and incubated for 15 s. Grids were blotted for 4 s at 4 °C with a blot force of 10 and plunge-frozen in liquid ethane cooled to LN2 temperature using a Vitrobot Mark IV (Thermo Fisher Scientific).

### Cryo-EM data acquisition

For data acquisition, frames were collected on a Titan Krios G3i transmission electron microscope (Thermo Fisher Scientific) equipped with a field emission gun, a BioQuantum post-column energy filter (Gatan), and a K3 direct electron detector (Gatan). Movies were recorded at an acceleration voltage of 300 kV in low-dose mode as dose-fractionated videos with a maximum image shift of 5 µm enabled by aberration-free image shift. In total, 16803 movies were collected, each with a total dose of 60.04 e^-^ /Å^2^ distributed over 31 fractions (1.94 e^-^ /Å^2^ per frame). Data were acquired in energy-filtered zero-loss mode (slit width 20 eV) in nanoprobe mode at a nominal magnification of 105kx, corresponding to a calibrated pixel size of 0.83 Å at the specimen level, using super-resolution mode and a 100 µm objective aperture. Defocus values ranged from ™1.0 to ™3.0 µm. Data were collected on Quantifoil R2/2 Cu 300 mesh grids, with 8 exposures acquired per hole.

### Cryo-EM image processing

The collected movies were corrected for beam-induced motion using patch alignment and dose-weighted with MotionCor2^24^. Contrast transfer function parameters were estimated with Gctf^25^. Particles were automatically picked using the Laplacian-of-Gaussian algorithm implemented in RELION 4^26^ and extracted with a pixel size of 1.10 Å. Subsequent 2D and 3D classifications were performed in RELION 4, and ribosomal particle subsets were imported into CryoSPARC v4.6.0^27^ for further refinement. Following an initial round of non-uniform refinement, an additional 3D classification step with a focused mask on the decoding and peptidyl transferase centers revealed two distinct conformational states (Supplementary Figure S2). These classes were refined independently by iterative non-uniform refinement and local refinement of individual subunits. Refinement was concluded once improvements in resolution and map quality became negligible. Final reconstructions were sharpened using standard B-factor correction in CryoSPARC. Local resolutions were estimated in cryoSPARC (FSC = 0.143) for the 80S ribosome and for the cleavage site using a mask encompassing the D6 region; histogram bins were computed from voxel-wise values within this mask.

### Model building and refinement

The previously published atomic model of the mouse 80S ribosome (PDB ID: 7CPU) was used as the initial template. The model was fitted into the cryo-EM density by rigid-body docking in UCSF ChimeraX. RNA and protein residues were removed in regions where the density was absent or highly fragmented. Structured elements were adjusted through iterative cycles of manual model building in Coot v0.9.8.95 ^28^ and real-space refinement in Phenix v1.21^29^, applying secondary structure restraints throughout. The mRNA and tRNAs were modeled based on the overall density features, as these represent an averaged signal from multiple heterogeneous conformations. The final models were validated using MolProbity, yielding favorable statistics for both states (Table S1).

### Assignment of rRNA modifications

Putative rRNA modifications were assigned by visual inspection of the cryo-EM density. Each nucleotide was examined for additional density features at the nucleobase and the 2′-OH group of the ribose sugar. Residues displaying such features were modeled as modified nucleotides in Coot. Given the local variation in map resolution, each potential modification site was evaluated individually by adjusting the map contour level to ensure consistency with the surrounding density.

### Statistical Analysis

No statistical methods were used to predetermine sample size. The cryo-EM data processing workflow, including particle numbers for each class, is detailed in Supplementary Figure S2.

### Figure generation

All figures showing structural models were generated using UCSF ChimeraX.

## DATA AVAILABILITY

Maps have been deposited to EMDB with accession codes EMD-55098 and EMD-55100.

## AUTHOR CONTRIBUTIONS

GL and MK designed the research, MG performed the experiments, analyzed the results, and drafted the manuscript; AR contributed to sample preparation; MK and CMTS contributed to data analysis. All authors contributed to the manuscript writing.

## ACKNOWLEDGEMENTS

We thank the Core Facility for cryo-Electron Microscopy (CFcryoEM) of the Charité-Universitätsmedizin Berlin for support in the acquisition (and analysis) of the data. The CFcryoEM was supported by the German Research Foundation (DFG) through grant No.

INST 335/588-1 FUGG Titan Krios 300 keV Kryo-Transmissions-Elektronenmikroskop. We thank Dr. Thiemo Sprink, Dr. Christoph Diebolder, Sabrina Golusik, and Metaxia Stavroulaki for their help with sample and grid preparation and data collection. We thank Dr. Julia Smirnova and Vasilii Mikirtumov for useful discussions. We thank Anika Mühlenberg, Gabriela Pflanz and Valérie Bégay for naked mole-rat animal care and colony management. The authors declare no competing interests.

