## Supplementary Material for "Cryo-EM structure of the naked mole-rat ribosome reveals a stabilized split 28S rRNA"

[5.] Charité – Universitätsmedizin Berlin, corporate member of Freie Universität Berlin and Humboldt Universität zu Berlin, Berlin, Germany

This file contains:

Figures S1, S2

Tables S1

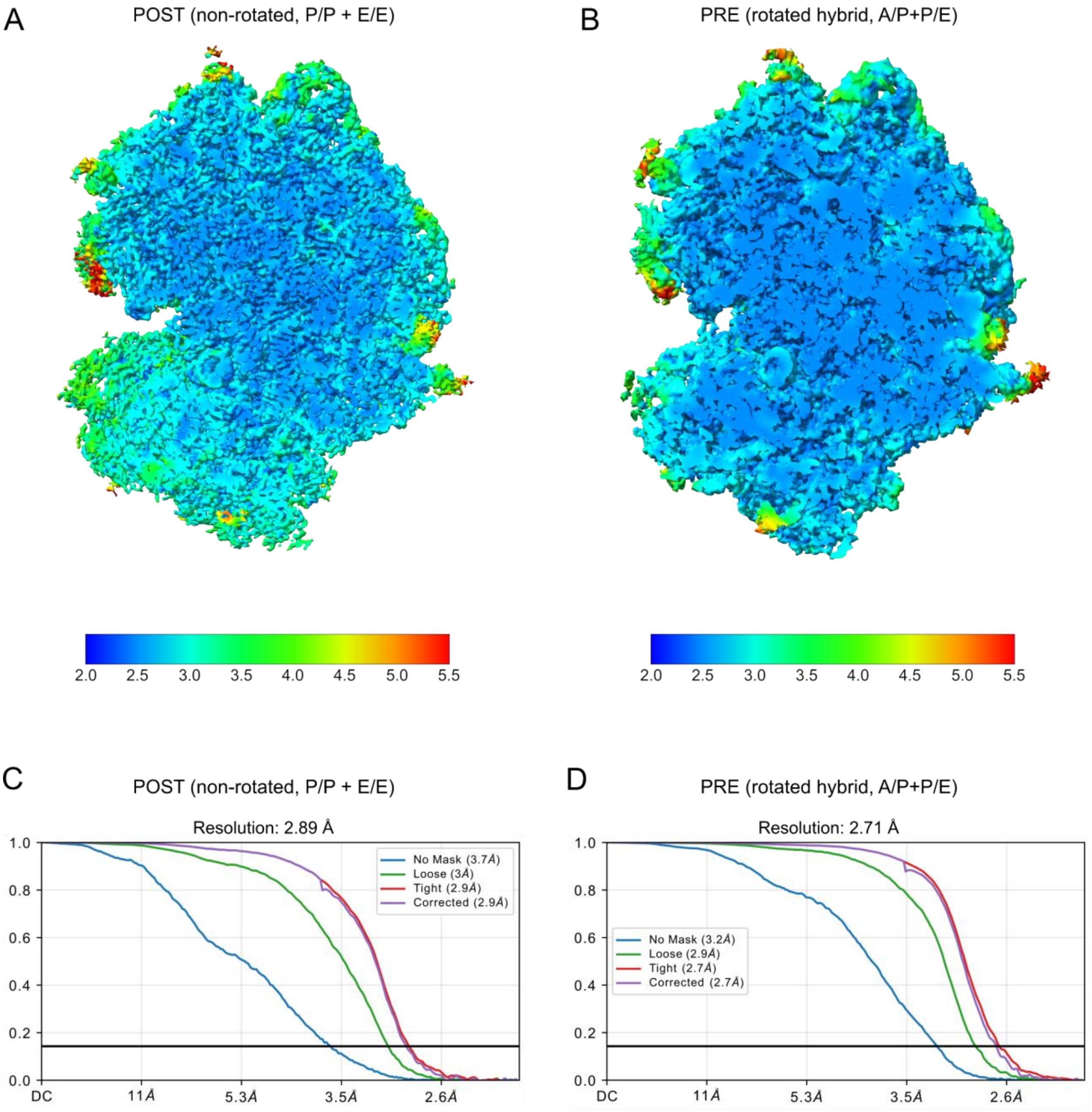


**Supplementary Figure S1.** Cutaway view through the cryo-EM density maps of the (A) POST and (B) PRE states. The maps are colored by the local resolution as calculated in cryoSPARC. GSFSC curves for the (C) POST and (D) PRE state structures. The resolution values are shown for the cutoff value of 0.143.

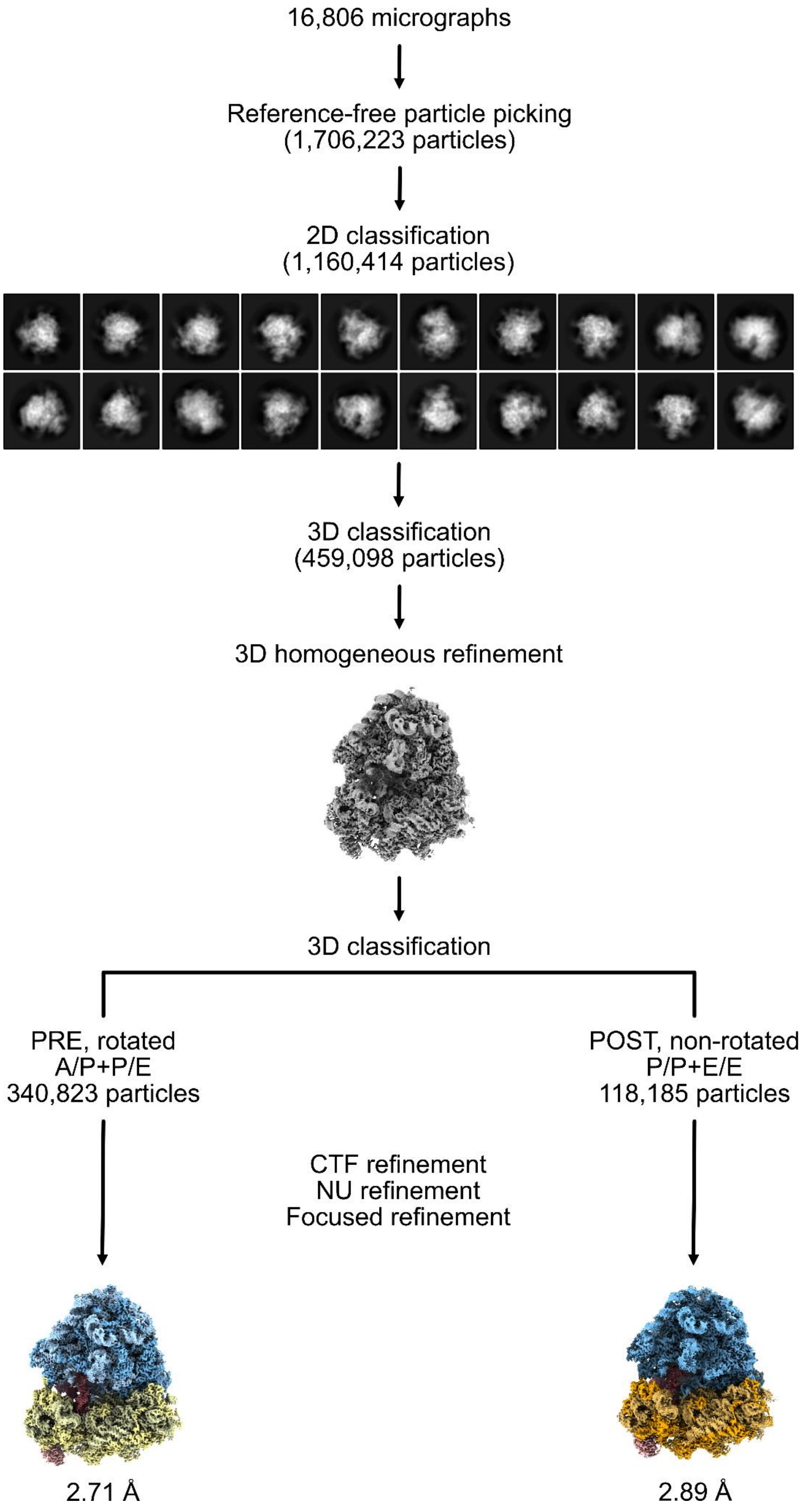


**Supplementary Figure S2.** Cryo-EM data processing workflow.

### Table S1: Cryo-EM Data Collection, Refinement, and Validation Statistics

| | NMR 80S (POST) | NMR 80S (PRE) |
|---|---|---|
| **Data Collection and Processing** | | |
| Microscope | Titan Krios G3i | Titan Krios G3i |
| Detector | Gatan K3 + Bioquantum GIF | Gatan K3+ Bioquantum GIF |
| Voltage (kV) | 300 | 300 |
| Electron dose (e−/A2) | 60.04 | 60.04 |
| Defocus range (µm) | -1.0 to -3.0 | -1.0 to -3.0 |
| Pixel size (Å) | 0.83 | 0.83 |
| Initial particles | 1706223 | 1706223 |
| Final particles | 118185 | 340823 |
| Map resolution (Å) | 2.9 | 2.7 |
| Map sharpening B-factor (A2) | -68.79 | -82.94 |
| **Model Refinement** | | |
| Initial model used (PDB ID) | 7CPU | 7CPU |
| Model components | 72 proteins, 4 rRNAs | 72 proteins, 4 rRNAs |
| Map-model FSC (FSC=0.5) | 3.05 | 2.83 |
| **Model Validation (MolProbity)** | | |
| MolProbity score | 2.53 | 1.79 |
| Clashscore | 12.97 | 4.72 |
| Ramachandran favored (%) | 88.74 | 90.20 |
| Ramachandran allowed (%) | 8.98 | 8.05 |
| Ramachandran outliers (%) | 2.29 | 1.75 |
| Rotamer outliers (%) | 2.71 | 0.96 |
| RMSD bonds (Å) | 0.008 | 0.003 |
| RMSD angles (°) | 1.070 | 0.777 |
